# Allele-specific antisense oligonucleotide treatment rescues *atad3-*associated phenotype in zebrafish

**DOI:** 10.64898/2026.05.20.726050

**Authors:** Shlomit Ezer, Shira Yanovsky-Dagan, Amit Granit, Matthew McDougal, Taeyoung Hwang, Israel Antman, Rotem Karni, Wan Hee Yoon, Ann Saada, Adi Inbal, Tamar Harel

**Author notes:** Corresponding author: Tamar Harel, M.D., Ph.D. Department of Genetics, Hadassah-Hebrew University Medical Center POB 12000, Jerusalem, Israel 9112001, Office.

## Abstract

Pathogenic variants in *ATAD3A* cause a spectrum of multisystem disorders, with a recurrent dominant-negative variant (c.1582C>T; p.Arg528Trp) associated with neurodevelopmental disease. Given the tolerance of *ATAD3A* to heterozygous loss of function variants, allele-specific transcript reduction represents a promising therapeutic strategy. We designed and optimized allele-specific antisense oligonucleotides (ASOs) targeting the c.1582C>T transcript and evaluated their efficacy and specificity in affected fibroblasts using allele-specific primers and amplicon-based next generation sequencing. Therapeutic potential was further assessed in vivo in zebrafish embryos expressing human wild-type or mutant *ATAD3A* transcripts. An optimized gapmer ASO selectively reduced mutant *ATAD3A* transcripts while relatively sparing the wild-type allele. In addition to RNase H-mediated degradation, the ASO induced exon skipping, leading to degradation of the aberrant transcript without production of a truncated protein. In zebrafish, expression of mutant human *ATAD3A* in embryos caused developmental abnormalities including reduced eye size, which were robustly rescued by co-injection of the optimized ASO. Our findings provide proof of concept for allele-targeted ASO therapy for dominant-negative *ATAD3A* variants. This work highlights the therapeutic potential of ASOs for rare dominant disorders involving genes tolerant to heterozygous loss-of-function, and establishes zebrafish as a versatile platform for in vivo ASO optimization.

## INTRODUCTION

*ATAD3A* is a nuclear-encoded gene encoding the mitochondrially localized AAA domain-containing ATPase 3A. Similar to other AAA+ proteins, it is predicted to function as a hexamer.^1^ Functionally, ATAD3A occupies a unique position within the mitochondrial architecture, spanning both the inner and outer mitochondrial membranes and thereby enabling diverse mitochondria-related activities. It has been implicated in the regulation of mitochondrial dynamics, as altered *ATAD3A* expression disrupts cristae organization and induces pronounced morphological changes, including mitochondrial fragmentation or elongation upon under- or over-expression.^2–4^ ATAD3A also facilitates mitochondrial DNA (mtDNA) maintenance by binding TFAM and regulating nucleoid trafficking and respiratory complex assembly. Beyond its roles in mitochondrial structure and genome stability, ATAD3A is essential for cholesterol metabolism, with loss of function leading to lipid accumulation and features of non-alcoholic fatty liver disease (NAFLD).^5^ Furthermore, it is a crucial component of mitochondria-associated membranes (MAM),^6^ where its interaction with the sigma-1 receptor and proper oligomerization is integral to maintain MAM integrity and to prevent neurological dysfunction.^7^ Additional studies have linked ATAD3A to innate immune responses via the cGAS-STING pathway^8^ and have identified it as an oncogenic factor that promotes chemoresistance in cancers through interactions with GRP78^9^ and regulation of PINK1-dependent mitophagy.^10^

Although pathogenic variants in *ATAD3A* are rare, their identification over the past decade has revealed a robust genotype-phenotype correlation. *ATAD3A*-associated diseases constitute a spectrum of multi-systemic disorders with predominant neurological features. At the severe end, biallelic loss of function (LoF) variants lead to severe brain malformations, respiratory insufficiency and neonatal lethality. At the milder end, a recurrent dominant *de novo* missense variant (NM_001170535.3(*ATAD3A*): c.1582C>T, p. (Arg528Trp)) causes developmental delay, optic atrophy, cardiomyopathy and peripheral neuropathy. The equivalent mutation in the *Drosophila* ortholog gene, *bor,* demonstrated a comparable phenotype and was shown to act in a dominant-negative (DN) manner.^11^

An emerging therapeutic approach for monogenic diseases, and more specifically in rare neurological diseases, is antisense oligonucleotide (ASO) treatment.^12,13^ Allele-specific ASOs are short synthetic oligonucleotides designed to selectively hybridize to pathogenic DNA or RNA transcripts and modulate gene expression in various ways, such as altered splicing or targeted RNA degradation.^14,15^

Gapmer ASOs represent a particularly effective subclass. They consist of a central DNA ‘gap’ flanked by chemically modified RNA nucleotides that hybridize to the target RNA. Formation of a DNA-RNA duplex recruits RNase H, leading to cleavage and degradation of the RNA.^16–18^ Chemical modifications can be added to the backbone or the bases to enhance nuclease resistance, binding affinity, specificity, and overall tolerability. Available modifications include 2’-O-methoxyethyl (2’MOE), the addition of a methoxyethyl group to the 2’ oxygen in the ribose; locked nucleic acid (LNA) sugars, with a methylene bridge connecting the 2’ oxygen and 4’ carbon in the ribose; or modifications to the backbone, such as a phosphorothioate (PS) backbone, replacing the oxygen in the phosphate group with a sulfur atom.^12,16,19^

In recent years, multiple research teams have successfully implemented ASO therapies relying on RNase H recruitment in preclinical animal models, with several advancing to FDA-approved status for DN or gain-of-function (GoF) monogenic disorders. In zebrafish, gapmer ASOs targeting developmental genes like *oep* and *smad2* achieved potent, allele-independent transcript knockdown via RNase H cleavage, validating the mechanism for sustained gene silencing beyond the limitations of morpholinos.^20^ Complementary rodent studies include SNP-specific gapmers that selectively reduce mutant *HTT* expression in Huntington’s disease models.^21^

Such preclinical successes underscore the versatility of ASOs for precision intervention in heterozygous dominant diseases. Tofersen, a gapmer ASO targeting *SOD1* mRNA for GoF-mediated amyotrophic lateral sclerosis (ALS), was approved by the FDA in 2023, after phase III trials demonstrated RNase H-mediated degradation of mutant transcripts, reduced neurofilament levels and slowed disease progression.^22–25^ For *SPTLC1*-associated childhood ALS driven by heterozygous GoF variants, allele-specific gapmer ASOs selectively knocked down mutant transcripts in vitro, normalizing enzymatic activity while sparing wild-type alleles.^26^ Preclinical examples include allele-specific gapmers for *KCNA2*-related drug-resistant epilepsy (heterozygous DN variant), which reduced seizure activity in neuronal cultures,^27^ and SNP-linked ASOs for Huntington’s disease, achieving selective mutant huntingtin lowering in mouse brain via intrathecal delivery.^21,28^ These RNase H-dependent strategies highlight the potential of ASOs for precision therapy in heterozygous dominant diseases.

We previously generated a zebrafish knockout model of the *ATAD3A* ortholog, *atad3*. Phenotypic characterization and transcriptomic analysis confirmed strong similarity between the human and zebrafish *ATAD3A*-related disorders, establishing zebrafish as a relevant in vivo model for mechanistic insight and therapeutic exploration.^29^ In this study, we use zebrafish to evaluate the therapeutic potential of ASOs targeting the recurrent dominant negative *ATAD3A* c.1582C>T variant. We show that ASO treatment effectively mitigates the phenotype induced by expression of the mutant human transcript in zebrafish, providing proof of concept for an ASO-based therapeutic strategy for *ATAD3A*-related disorders.

## RESULTS

### ASO screening for maximal efficiency and specificity

To identify the most effective ASO, we first designed ten gapmers (ASO1-10) tiled across the NM_001170535.3(*ATAD3A*): c.1582C>T variant (**Supplemental Table 2)**. An in silico off-target search in GGGnome confirmed that each candidate was at least two mismatches away from any other coding region. For initial screening, heterozygous fibroblasts from an affected individual were transfected with 100nM of each ASO for 48hr, and the expression levels of total *ATAD3A* and of the mutant allele were compared (**Fig. 1A**), using different sets of primers. Gene expression was normalized to *ATAD3A* expression in untreated fibroblasts. Among the initial candidates, ASO4 showed the highest efficiency, reducing the mutant allele by 91%. However, the reduction was not specific for the mutant allele as it reduced the total gene expression by 84%. To improve specificity, we generated six modified versions of ASO4 containing one or two additional mismatches (ASO4a-f). A scrambled ASO control was designed based on ASO4a (**Supplemental Table 2)**, and was confirmed in silico not to have other targets. All added mismatches increased allele specificity, with ASO4a and ASO4f showing the most favorable balance of efficacy and specificity (**Fig. 1B, Supplemental Fig. 1).**

**Figure 1:**
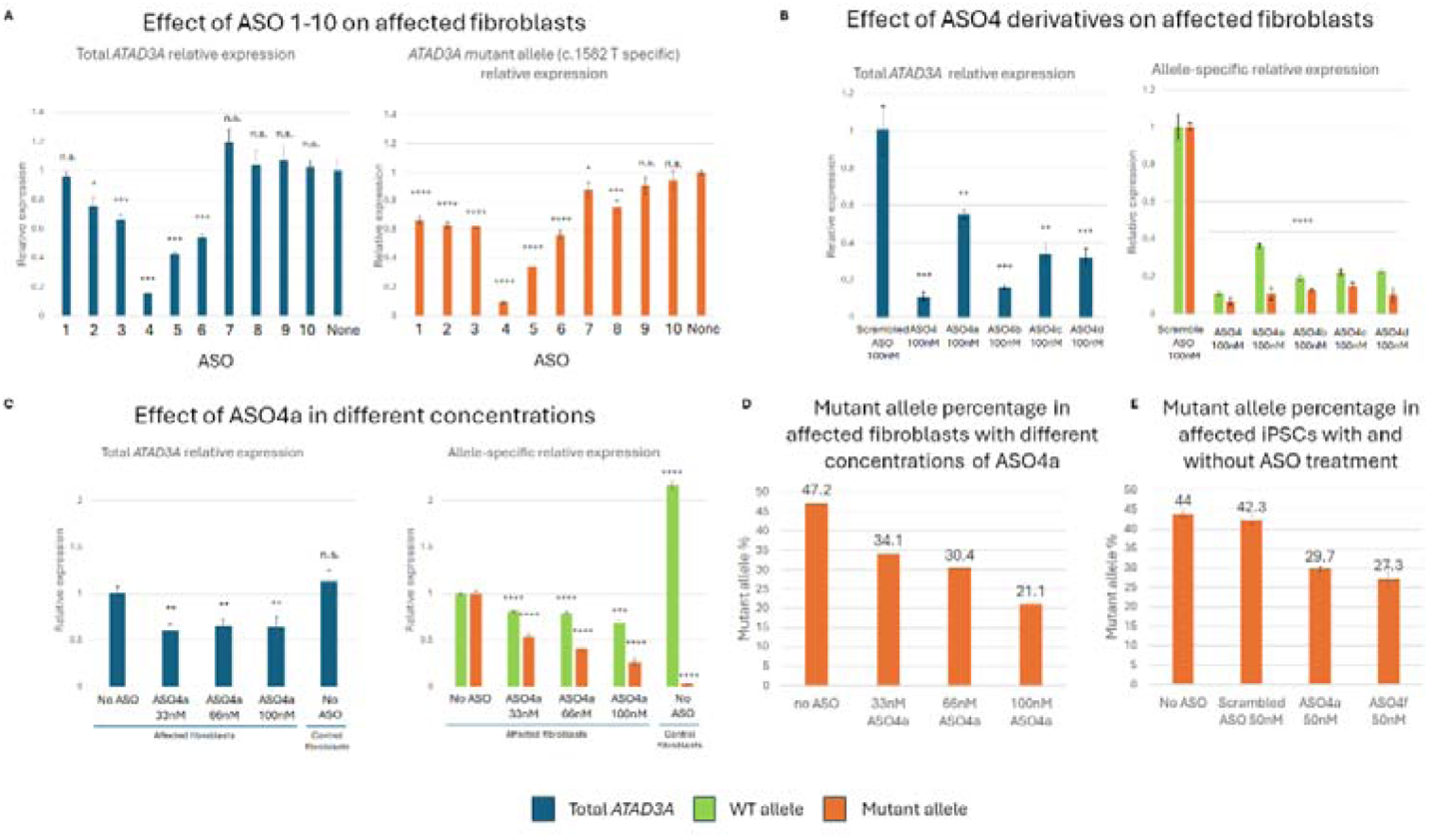
ASO trials on heterozygous fibroblasts. **(A)** Total *ATAD3A* and mutant RNA expression in affected fibroblasts treated for 48h with 100nM ASO (1-10), compared to untreated cells. Asterisks above each column represent level of significance compared to untreated affected cells (*p val<0.05, **p val<0.01, ***p val<0.001, ****p val<0.0001, n.s. not significant). **(B)** Total and allele-specific *ATAD3A* expression in affected fibroblasts treated with 100nM of ASO4, ASO4a-d compared to cells treated with scrambled ASO. Asterisks above each column represent level of significance compared to affected cells treated with scrambled ASO. **(C)** Total and allele-specific *ATAD3A* expression in affected fibroblasts treated with ASO4a in different concentrations (33nM, 66nM, and 100 nM). Asterisks above each column represent level of significance compared to untreated affected cells. **(D)** Percentage of mutant allele in the RNA of affected fibroblasts treated with different concentrations of ASO4a for 48h determined by Ampliseq. **(E)** Percentage of mutant allele from cDNA of affected patient-derived iPSCs treated with 50nM of ASO4a and ASO4f for 30h. Allele percentage was calculated by PCR followed by ONT sequencing. The allele frequencies were calculated from the relative fractions of C (WT) and T (mutant) alleles in the control cell line, and were normalized to expression in the scrambled ASO treatment.

ASO4a showed consistent, dose-dependent allele-specific effects by both allele-specific RT-qPCR and amplicon-based next generation sequencing (Ampliseq). By allele-specific RT-qPCR, the highest tested concentration of ASO4a (100nM) reduced the mutant allele by 74% and the WT allele by 32% relative to untreated affected fibroblasts (**Fig. 1C),** corresponding to a ∼two-fold preferential reduction of the mutant allele. We reasoned that this might be sufficient to alter the stoichiometry of the encoded ATAD3A subunits within the hexamer. Quantification of the relative mutant allele fraction among total transcripts as measured by Ampliseq, showed a 55% reduction in the mutant allele at the same ASO concentration (**Fig. 1D).**

To further evaluate the optimized ASOs, ASO4a and ASO4f, in an independent cellular model, we utilized an iPSC line originally derived from the same patient, together with an isogenic control line (T. Bae et al., manuscript submitted). In patient-derived iPSCs, a concentration of 50nM was utilized to minimize ASO-mediated cell death. ONT sequencing spanning the c.1582 variant region was used to assess the mutant to WT allele ratio. At 50nM, both ASOs reduced the mutant allele frequency compared to the scrambled ASO control: ASO4a by 29.8% and ASO4f by 35.5% (**Fig. 1E**). Baseline ONT sequencing showed a modest allelic imbalance, with slightly higher expression of the C (WT) allele compared to the T (mutant) allele (C:T = 56:44) (**Fig. 1E**); all allele-specific changes were calculated relative to this baseline ratio. Together, our data from multiple cellular models suggest that targeting mutant *ATAD3A* with ASOs may have therapeutic potential.

### ASO4 and its derivatives cause exon skipping leading to transcript degradation

RNA from heterozygous cells treated with ASO4 and 4a-f showed an additional peak with a lower melting temperature in the melt-curve analysis of the RT-qPCR for the exon 14-16 product (**Fig. 2A**). A faint band shorter than the exon 14-16 product was also seen on agarose gel, increasing in intensity with the increasing ASO concentration, and more prominent in affected cells than in WT controls (**Fig. 2**B). In WT fibroblasts, ImageJ quantification revealed an 11% increase in the shorter product at the high ASO4a concentration (100nM) compared to its scrambled ASO treatment control. In affected fibroblasts, the parallel relative increase in the exon-skipping product was 420% (**Fig. 2C-D**). Sequencing of the shorter product revealed skipping of exon 15 (containing the c.1582 variant) (**Fig. 2E**).

**Figure 2:**
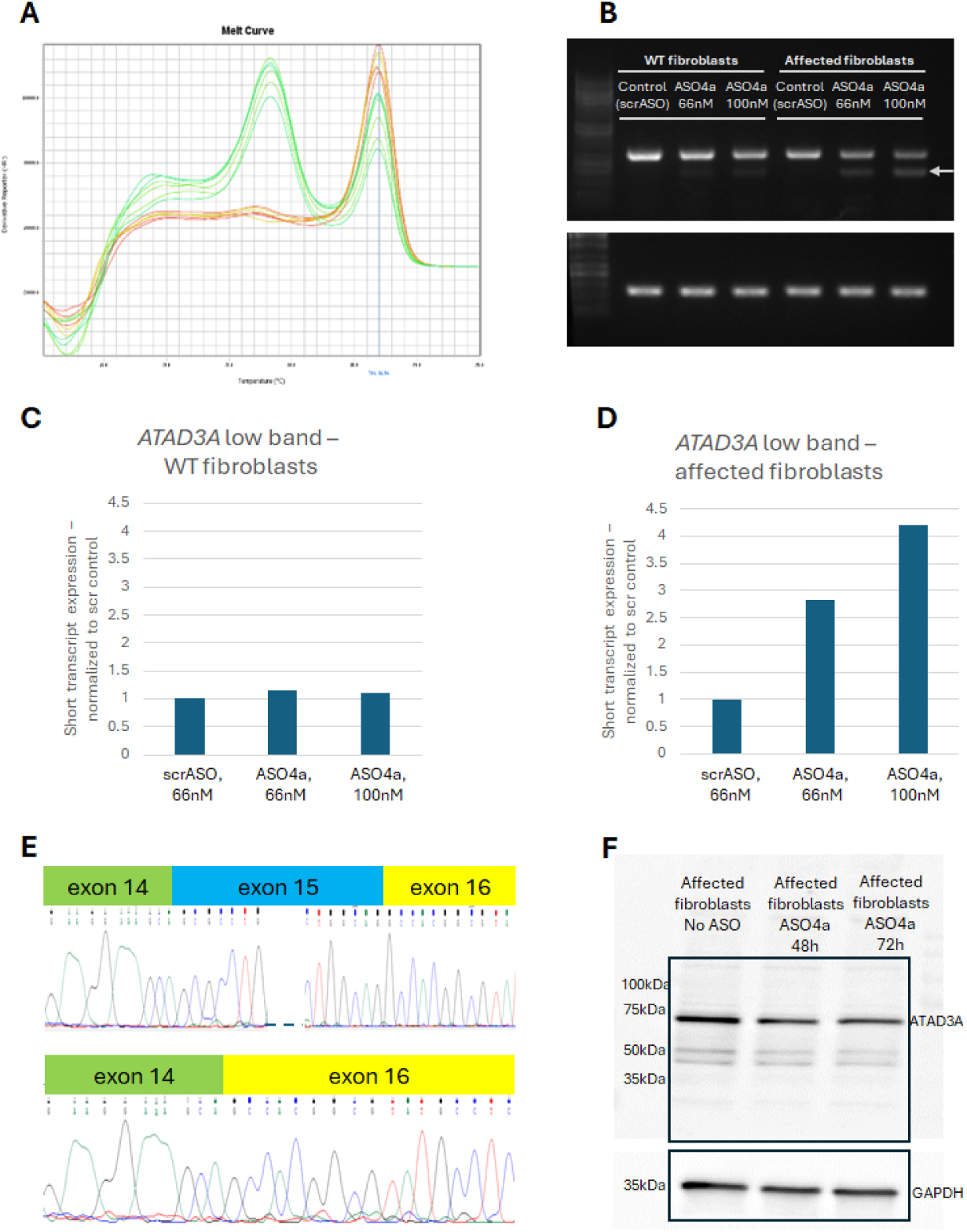
ASO4 and its derivatives cause exon skipping. **(A)** RT-qPCR on affected fibroblasts treated with ASO4a show two peaks for exon 14-16 primers. In red, the melt-curve of untreated cells with a single peak. In green, two peaks: the left one (lower melting temperature) indicating a shorter product. **(B)** Exon 14-16 RT-PCR products show an additional shorter product on agarose gel (marked by an arrow), correlating in intensity with ASO4a concentration, more so in unaffected fibroblasts compared to WT fibroblasts. Lower panel is RPLP0 housekeeping gene for quantification. **(C-D)** Quantification of the shorter *ATAD3A* band. Each sample is normalized to its RPLP0 expression (housekeeping gene) and to its scrambled ASO treated control (66nM); **(C)** WT fibroblasts, **(D)** affected fibroblasts. **(E)** Sanger sequencing chromatograms of RT-PCR exon 14-16 products after ASO4 treatment. Upper panel: main product band, with the expected exon 14-16 sequence. Lower panel: faint lower band indicating the skipping of exon 15. **(F)** Western blot of affected fibroblasts lysate with ASO4a (100nM) and exon-skipping shows no shorter protein product compared to untreated cells. Upper panel: anti-ATAD3A; lower panel: anti-GAPDH.

Analysis by the ESEfinder tool^30,31^ predicted that several exonic splicing enhancer (ESE) motifs are located within the target sequences of ASOs 1-10. Specifically, the targeting site of ASO4 and its derivatives (ASO4a-f) includes predicted binding sites for SRSF1 (Serine and Arginine Rich Splicing Factor 1) and SRSF6 (**Supplemental Fig. 2A**).

Exon 15 is out-of-frame (109bp long), and its skipping results in a premature stop codon in exon 16, suggesting that the transcripts lacking exon 15 may trigger nonsense mediated decay (NMD) or otherwise be translated into an unstable protein (**Supplemental Fig. 2B)**. To determine whether a stable C-terminally truncated protein was produced by the aberrant transcript, lysates from heterozygous fibroblasts treated with ASO4a for 48 or 72h were analyzed by Western blot for total ATAD3A. Lysates for GAPDH detection was loaded in different wells, to avoid obscuring detection of a smaller truncated product. Although RNA from the treated cells showed marked exon skipping (**Fig. 2A-B,D**), no shorter protein was detected compared to untreated fibroblasts (**Fig. 2F**), indicating that exon skipping led to degradation of the mis-spliced allele at either the mRNA or protein level. Together, these findings suggest that the efficacy of the ASOs can be attributed to both RNase H-dependent cleavage and induction of exon skipping.

### WT *ATAD3A* rescues the *atad3*-null phenotype in zebrafish embryos

Due to the high protein homology between human and zebrafish, we hypothesized that the human WT *ATAD3A* could rescue the phenotype of embryos lacking the endogenous ortholog gene. To test this hypothesis, human WT *ATAD3A* cDNA was cloned into PCS2+ plasmids. Following site-directed mutagenesis, both the WT and mutant copies were transcribed and isolated as RNA. Sequencing of the plasmids confirmed WT and mutant genotypes (**Supplemental Fig. 3**). *Atad3*-KO heterozygous fish^29^ were crossed, and embryos were injected with 250pg of either WT or mutant human *ATAD3A* RNA. At 3dpf, embryos were imaged and then genotyped, to allow measuring of eye size specifically for the *atad3*-null embryos (injected or un-injected). Un-injected *atad3-*null mutants had an average eye size 27% smaller than that of un-injected unaffected siblings (**Supplemental Fig. 4)**. Injection of WT human *ATAD3A* rescued the reduced-eye-size phenotype, with WT-*ATAD3A-*injected embryos showing an average eye size 15% larger than that of un-injected *atad3-*null embryos. In contrast, injection of mutant human *ATAD3A* RNA worsened the phenotype, with mutant-*ATAD3A*-injected embryos showing an average eye size 17% smaller than that of un-injected null embryos. Together, these results demonstrate functional complementation of the orthologous zebrafish gene by human WT *ATAD3A* and suggest that mutant *ATAD3A* has a deleterious effect in vivo.

### Injection of mutant human *ATAD3A* RNA, but not WT RNA, into zebrafish embryos causes a phenotype similar to *atad3-*null embryos

To investigate the ASO effect in vivo, we first expressed human *ATAD3A* RNA in wild-type AB/TL zebrafish embryos to characterize the effect of WT and mutant human *ATAD3A* in this model. This experiment was designed to ensure that human WT *ATAD3A* is not toxic to zebrafish embryos, and to determine whether mutant human *ATAD3A* induces a measurable phenotype. Wild-type *ATAD3A* AB/TL zebrafish embryos were microinjected with 125, 250 or 375pg of either WT or mutant human *ATAD3A* RNA. The WT RNA caused no significant change in eye size compared to un-injected embryos in all concentrations tested (**Fig. 3A)**, indicating that the human *ATAD3A* RNA does not have a dominant-negative effect on endogenous zebrafish Atad3, nor a toxic gain-of-function effect. However, injection of the mutant RNA caused a significant reduction in eye size (p val < 0.01) (**Fig. 3B**), ranging from 23% to 39% reduction compared to un-injected embryos. Although there was no significant difference between the concentrations, the number of under-developed embryos whose eyes could not be measured was positively correlated with the amount of injected mutant RNA. Numbers of embryos imaged and measured are provided in **Supplemental Table 3.** The difference in eye size can be seen in representative images of embryos injected with 250pg WT vs. mutant RNA (**Fig. 3C-D** respectively). Additional data from embryos injected with different concentrations of WT RNA showing minimal change in eye size are presented in **Supplemental Fig. 5.**

**Figure 3:**
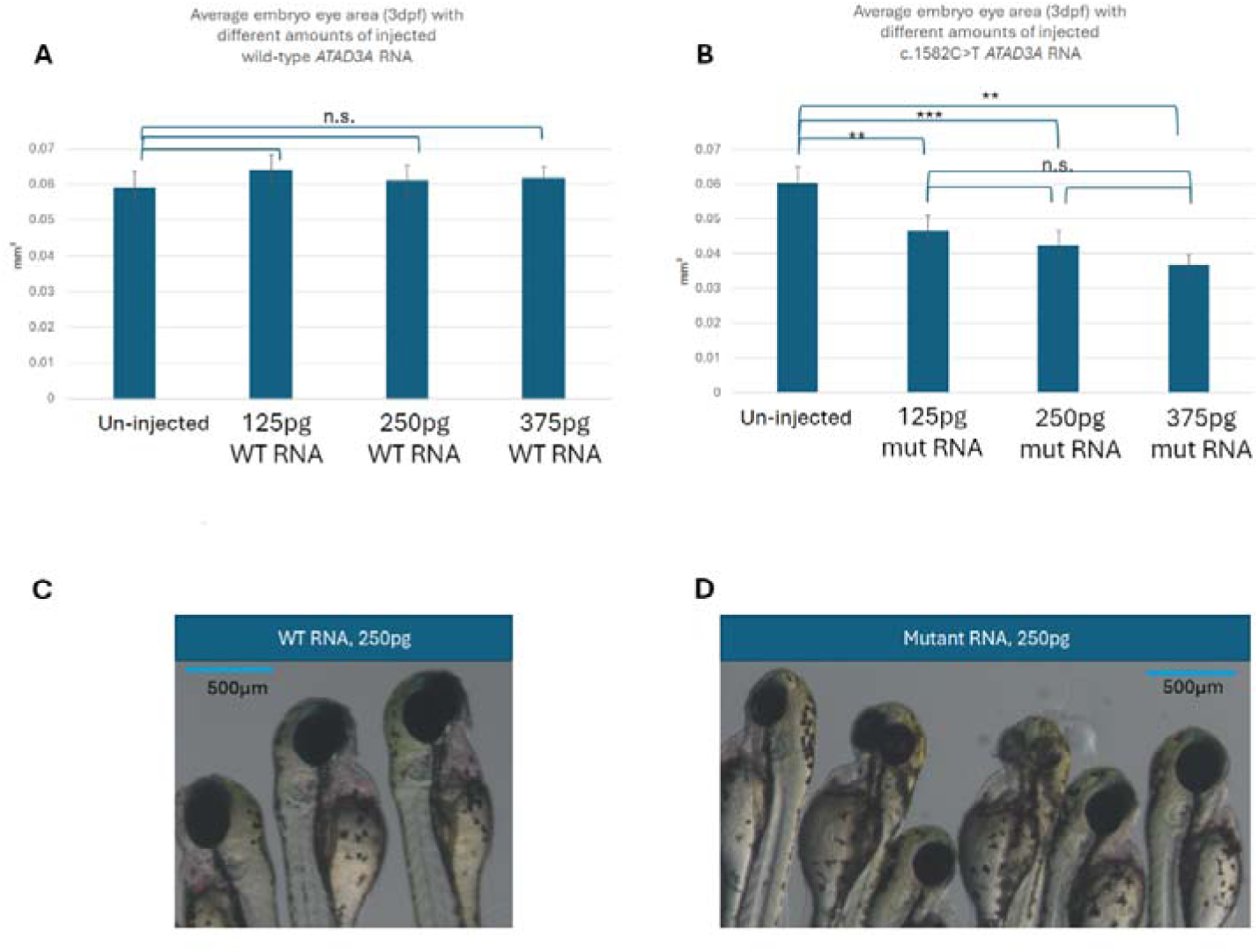
Mutant human *ATAD3A* RNA causes aberrant phenotype in zebrafish embryos. **(A)** Injection of WT *ATAD3A* RNA did not affect eye size at 3dpf at 125pg, 250pg or 375pg compared to un-injected embryos. (**B**) Injection of c.1582 C>T mutant *ATAD3A* RNA caused a reduction in eye size at 3dpf in all three amounts injected. (**C**) Representative image of WT-injected embryos at 3dpf, showing larger head and eye sizes compared to (**D**) a representative image of mutant-injected embryos (250pg RNA). Asterisks represent levels of significance (**p val<0.01, ***p val<0.001, n.s. not significant).

### ASO4a rescues the phenotype caused by microinjection of the mutant *ATAD3A*

The effects of mutant RNA injected into AB/TL embryos – namely, reduced eye size, reduced head size and increased cardiac edema **-** were reversed by the subsequent addition of ASO4a, but not by scrambled ASO (**Fig. 4A-C, Supplemental Fig. 6A-B**). Injection of the ASO alone (either 4a or scrambled) had no significant effect on eye, head or heart size compared to un-injected embryos. Numbers of embryos imaged and measured are provided in **Supplemental Table 3**. Embryos injected with RNA, with or without ASO, had both eyes measured as some embryos had variability between eyes.

**Figure 4:**
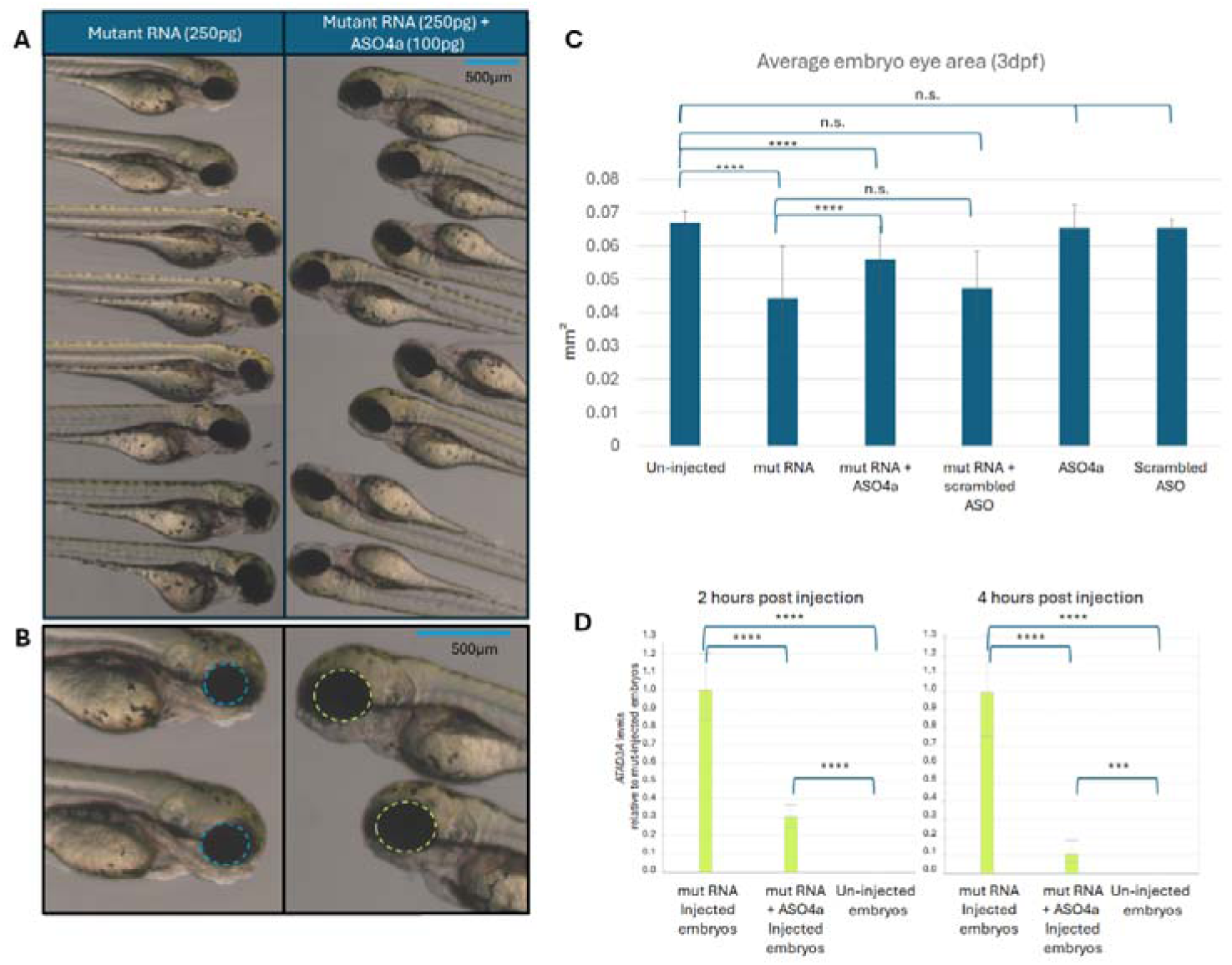
Injection of ASO4a following mutant RNA ameliorates phenotype. **(A)** Representative images of 3dpf embryos injected with either mutant RNA (on the left) or mutant RNA followed by ASO4a (on the right). **(B)** Close-up image showing larger eye size in embryos injected with ASO4a following mutant RNA injection. **(C)** Average eye size at 3dpf. Embryos are either un-injected or injected with RNA (250pg) and/or ASO (100pg), as indicated. **(D)** Human-*ATAD3A* RNA levels in embryos injected with mutant RNA only (250pg) compared to embryos injected with mutant RNA (250pg) followed by ASO4a (100pg). RNA extraction 2h post injection on the left, 4h post injection on the right. Asterisks represent levels of significance (***p val<0.001, ****p val<0.0001, n.s. not significant).

The ASO effect on the mutant RNA was also tested at the RNA level. Two hours post injection, the human *ATAD3A* RNA level in the mutRNA+ASO4a group was 70% lower than in the mutRNA without ASO group (pools of 13 embryos); four hours post injection, human *ATAD3A* RNA level in the mutRNA+ASO4a group was 90% lower than in the mutRNA without ASO group (pools of 11 embryos). No human *ATAD3A* was detected in un-injected embryos (**Fig. 4D**). This supports our assumption that the ASO targets and degrades the injected mRNA.

## DISCUSSION

In this study, we aimed to reduce expression of the recurrent dominant-negative *ATAD3A* c.1582C>T variant, in order to shift the balance of ATAD3A hexamers towards wild-type subunits. We designed and optimized allele-targeted ASOs, quantified their total and allele-specific effects in affected fibroblasts, and refined specificity by introduction additional mismatches. Their effect was subsequently replicated in iPSCs. The lead ASO was then evaluated in a zebrafish embryo model after confirming that expression of human wild-type *ATAD3A*was not detrimental in this system.

We hypothesized that an allele-specific ASO would be a promising therapeutic strategy for the *ATAD3A* c.1582 C>T variant based on the genetic characteristics of this gene. *ATAD3A-*associated disease causes both dominant and recessive inheritance, depending on the variant, and individuals heterozygous for LoF variants are clinically unaffected, indicating tolerance to reduced gene dosage. This concept is broadly applicable to genes in which pathogenicity arises from monoallelic GoF or dominant-negative variants, as well as from biallelic (but not monoallelic) LoF, since selective reduction of the mutant allele is expected to be tolerated.^32^

Our data suggest that ASO4 and its derivatives reduce mutant *ATAD3A* expression through two complementary mechanisms. First, as expected for gapmer ASOs, hybridization of the ASO to the target transcript recruits RNase H to the DNA-RNA hybrid, resulting in degradation of the mutant mRNA.^16^ Second, we found that ASO treatment affected splicing and induced skipping of exon 15, which contains the pathogenic variant. We did not detect a truncated ATAD3A protein after ASO treatment, indicating that this transcript most likely underwent nonsense mediated decay or encoded an unstable protein. The predicted presence of an ESE motif within the ASO4 target region suggests that exon skipping may result from disruption of splicing regulatory elements.^30,31^ This additional mechanism may be advantageous, provided that ASO hybridization remains sufficiently allele-selective. Unlike classical splice-switching oligonucleotides (SSOs), which are typically designed to restore normal splicing or bypass exons with pathogenic LoF variants in order to restore the normal reading frame,^14,33,34^ our data suggest that ASOs can also be used to induce mis-splicing as a means of eliminating an aberrant transcript. Notably, the zebrafish model isolates RNAse H-dependent activity, since the injected RNA is already spliced and therefore bypasses any ASO-induced splice modulation.

The ASO-induced reduction in ATAD3A protein levels was less pronounced than the reduction in RNA level. Although technical factors may contribute to this difference, it may also reflect the greater stability of the protein relative to the transcript. Accordingly, protein measurements may provide a more conservative estimate of the biological effect.

The main achievement of this work is proof of concept of the phenotypic rescue in vivo. These results support the therapeutic potential of our ASO, calling for continued optimization of ASO specificity and potency. Further studies across additional cellular models should help clarify the mechanisms underlying exon skipping and refine the balance between efficacy and selectivity.

Several considerations need to be taken into account when designing ASO treatment. Off-target activity remains a central concern, as partial complementarity to unintended transcripts can lead to unanticipated gene knockdown or splice modulation, particularly with high-affinity chemistries such as phosphorothioate backbones and locked nucleic acids.^35^ We reduced this risk by in silico off-target screening using GGGenome (https://gggenome.dbcls.jp/), although the most likely off-target effect in this setting remains partial hybridization to the WT allele. Emerging strategies to enhance allele specificity include exploiting differences in RNA secondary structure between mutant and wild-type transcripts and alternating DNA and RNA nucleotides within the ASO to favor binding to structurally accessible mutant regions.^36^ Incorporation of an additional mismatch complementary to the open region in the secondary structure increased specificity 10-fold in a particular study.^35^

Delivery presents an additional challenge. ASOs must traverse biological barriers, enter target tissues and cells, and escape endosomal compartments. This often calls for invasive routes such as intrathecal (IT) administration for targets in the central nervous system (CNS), or other specialized delivery vehicles, yet introduced procedural risks and manufacturing challenges.^37^ While zebrafish embryos allow for yolk-sac microinjection, translation to human therapy will require careful optimization of delivery routes, dosing schedules, and long-term safety. Finally, ASO stability is a double-edged sword: insufficient nuclease resistance leads to rapid degradation and loss of efficacy, whereas extensive chemical modification to enhance stability can increase protein binding, alter pharmacokinetics, trigger immune activation, and narrow the therapeutic window.^37^ Further studies comparing alternative chemistries and modification patterns will be required for clinical translation of *ATAD3A*-targeting ASOs.

An additional limitation of the current study is that allele specificity was rigorously assessed in vitro but not fully evaluated in vivo. Zebrafish experiments involved injection of either wild-type or mutant human RNA individually but not co-expression of both alleles. Consequently, while our results clearly demonstrate in vivo efficacy, future studies incorporating co-injection of wild-type and mutant RNA should enable more direct evaluation of allele specificity in vivo. Complementary validation in additional cellular systems, including iPSC-based models such as neurons or organoids, will further strengthen translational relevance.

Importantly, complete allele specificity may not be necessary for therapeutic benefit. Because ATAD3A acts as a homomeric hexamer, even partial preferential reduction of the mutant transcript could substantially increase the proportion of fully functional complexes. In heterozygous cells, many hexamers are expected to incorporate at least one mutant subunit, compromising function of the entire protein complex. Accordingly, shifting allelic stoichiometry toward the wild-type transcript, even without full preservation of wild-type expression, may therefore produce meaningful biological benefit. This suggests that careful dose titration to achieve partial, allele-biased knockdown may represent a viable therapeutic strategy.

In conclusion, our study provides proof of concept that an ASO targeting the recurrent monoallelic c.1582C>T variant can efficiently degrade the mutant transcript in vivo and rescue disease-relevant phenotypes in a zebrafish model. These findings support effective targeting of pathogenic RNA and establish zebrafish as a useful platform for in vivo ASO optimization. More broadly, this work demonstrates the therapeutic potential of ASO-based approaches for rare dominant disorders involving genes tolerant to heterozygous loss of function, and supports the feasibility of precision, variant-specific interventions, including future N-of-1 strategies.^38–42^

## MATERIALS AND METHODS

### Cell maintenance

Fibroblasts: Fibroblasts from an affected individual (reported in Harel at el. 2016 and consented under research protocol H-29697) were maintained in DMEM Dulbeccòs Modified Eagle Media high glucose supplemented with 15% fetal bovine serum, 2mM L-glutamine and 1% penicillin-streptomycin (Biological Industries, Beit Haemek, Israel).

Induced Pluripotent Stem Cells (iPSCs): iPSCs were generated in accordance with OMRF IRB 26-05, and were grown on Matrigel (Corning 354277) coated 6-well plates in StemFlex medium (Gibco A3349401) and regularly maintained by clump passaging using ReLeSR (StemCell Technologies # 100-0484).

### ASO design and preparation

ASOs were designed spanning the *ATAD3A* c.1582 C>T variant. GGGenome (https://gggenome.dbcls.jp/) was used for detection of off-target sequences. 2’MOE gapmers were ordered from BioSpring (Germany) and IDT (USA). Gapmers were composed of 5 modified bases (in BioSpring: 2’MOE-5Me-rU, 2’MOE-rA, 2’MOE-5Me-rC, 2’MOE-rG; in IDT: 2’MOE-A,G,T or MeC), 10 DNA bases, and 5 modified RNA bases, with phosphorothioate bonds.

### Transfection

Fibroblasts: The day before transfection, 10^5 fibroblasts per a 6-well plate well were plated. They were next transfected using either Lipofectamine CRISPRMAX (Thermo Fisher, USA) or Lipofectamine RNAiMAX (Thermo Fisher, USA) with different concentrations of various ASOs for 48 hours, prior to collection in TRIzol Reagent (Thermo Fisher, USA) for RNA extraction.

iPSCs: The day before transfection, cells were dissociated using Accutase (StemCell Technologies # 07920) and 2-2.5×10^6 cells per well were plated in StemFlex medium containing 10µM Y-27632 dihydrochloride ROCK inhibitor (StemCell Technologies #72304) on Matrigel coated 6-well plates. As StemFlex media may inhibit transfection, iPSCs were changed to mTESR1 medium (StemCell Technologies #85850) containing 10µM Y-27632 dihydrochloride ROCK inhibitor to increase cell viability immediately before transfection. Formed transfection complexes utilizing Lipofectamine Stem Transfection Reagent (Invitrogen STEM00008) containing 50nM final concentration of ASO were then delivered to cells. 24 hours post-transfection, media was changed to 1:1 mix of mTESR1:StemFlex media. After another 6 hours of incubation (30h from transfection), cells were harvested with TRIzol reagent (Invitrogen 15596026) and samples were stored at −80°C until RNA isolation. RNA was isolated using the Zymo RNA Clean & Concentrator-25 kit (Zymo R1017) following manufacturer instructions and stored at −80°C until cDNA synthesis.

### RNA extraction and cDNA synthesis

Fibroblasts: RNA was extracted from cells using TRIzol Reagent (Thermo Fisher, USA). cDNA was prepared using the qScript cDNA Synthesis Kit (Quantabio, USA). cDNA from embryos injected with human RNA was synthesized using qScript Flex cDNA Synthesis Kit (Quantabio, USA) with random primers.

iPSCs: cDNA synthesis was performed using the SuperScript IV First-Strand Synthesis System (Invitrogen 18091050) using oligo dT primer following manufacturer instructions.

### Reverse Transcription-quantitative Polymerase Chain Reaction (RT-qPCR)

Relative gene expression was assessed by RT-qPCR using PerfeCTa SYBR Green FastMix (Quantabio, USA). Primers are listed in **Supplemental Table 1**.

### Amplicon-based next generation sequencing (Ampliseq)

Primers complementary to regions flanking the c.1582 C>T variant, attached with adapters for Ampliseq PCR reaction, were designed and ordered from IDT (USA) (**Supplemental Table 1)**. PCR for the amplification of this region with the addition of adapters was performed on cDNA from ASO treated fibroblasts using PCRBIO Taq Mix Red. An additional PCR was performed to attach sample-specific index sequences (IDT, USA) to the adapter. PCR products were purified using KAPA Pure Beads (Roche, Switzerland) and diluted to a normalized concentration of 0.1ng/ul. Samples were sequenced on the NovaSeqX platform (Illumina) using a 161 paired-end sequencing scheme, resulting in 1-2M reads per sample. cDNA AmpliSeq analysis was performed using the following pipeline: sequenced reads were aligned to the reference genome using STAR, with a genome index built for 150 bp reads. The resulting BAM files were post-processed to ensure compatibility with GATK requirements. Variant calling was performed using GATK Mutect2 to estimate the allele frequency (AF) of each allele in each sample. In addition, GATK AlleleCount was used to compute the raw allele frequencies for each sample.

### Long-read sequencing of amplicon (ONT sequencing)

Following cDNA synthesis, a subsequent PCR to amplify *ATAD3A* was performed using the Q5 High-Fidelity DNA Polymerase (New Englad BioLabs M0491S) following manufacturer instructions from 200ng total cDNA. PCR products were purified using the QIAquick PCR Purification Kit (Qiagen 28106). Primers are listed in **Supplemental Table 1**.

Raw sequencing reads were aligned to the reference genome (hg38) using minimap2 (v2.30)^43^ with the splice-aware option enabled.

### Western blo**t**

Protein was extracted from fibroblast pellets using RIPA and protein concentration was detected using Pierce™ Bradford Protein Assay Kit (Thermo Fisher, USA). For ATAD3A detection, 25ug of protein of each sample were loaded onto a 4-20% gel (BioRad, USA). For GAPDH detection, 10ug were loaded. Following transfer, blocking of the membrane was performed in 5% milk in Tris-Buffered Saline with Tween 20 (TBST). ATAD3A was detected using H00055210-D01 antibody (NOVUS Biologicals, USA), GAPDH was detected using ab8245 antibody (Abcam, UK). Secondary antibodies used were ab6721 (anti rabbit) and ab6789 (anti mouse), (Abcam, UK). All antibodies were used in 5% BSA in TBST.

### Zebrafish maintenance

We used AB/TL hybrid zebrafish (*Danio rerio*) as wild-type (WT), and a CRISPR/Cas9-generated heterozygous line for an *atad3* deletion.^29^ Adults were maintained according to standard procedures on a 14-h light/10-h dark cycle at 28°C. Embryos were produced by pair mating and raised at 28.5°C in egg water (0.3 gr NaCl in 1-liter distilled H_2_O). *atad3*^+/-^ zebrafish were kept on an AB/TL WT background. Guide for the Care and Use of Laboratory Animals guidelines were followed regarding animal maintenance and throughout all experiments presented in this work. The research was approved by the animal ethics committee of the Hebrew University of Jerusalem (MD-20-16420-1).

### Construction of plasmid and mRNA synthesis

The PCS2+ plasmid with an NM_001170535.3(*ATAD3A*) insert was generated using Gibson assembly. Briefly, the same pair of primers was used to perform PCR for the linearization of an empty PSC2+ plasmid and to amplify an *ATAD3A* insert previously cloned into a pEZ-Lv105 plasmid. Original DNA templates were digested using DpnI, and PCR products were purified from agarose gel electrophoresis. Gibson assembly was performed with the linearized PSC2+ plasmid and the *ATAD3A* insert according to standard protocol.^44^

The assembled plasmid was transformed into DH5α competent *E. coli* cells and amplified using the NucleoBond Xtra Midi kit (Macherey-nagel, Germany). Site-directed mutagenesis was performed using the Q5 Site-Directed Mutagenesis Kit (New England BioLabs, USA) using back-to-back mutagenesis primers (listed in **Supplemental Table 1**) followed by plasmid purification, transformation, and midi-scale amplification as described.

For in vitro transcription, plasmids were linearized with the NotI-HF restriction enzyme (New England Biolabs, USA), excised from gel and cleaned using the NucleoSpin Gel and PCR clean-up kit (Macherey-nagel, Germany). Capped mRNA was synthesized using the SP6 mMESSAGE mMACHINE kit (Invitrogen, USA) according to manufacturer’s instructions, and purified using the GenElute–E single spin RNA cleanup kit (Sigma-Aldrich, USA).

### Microinjections, phenotyping and genotyping of zebrafish embryos

Wild-type or mutant *ATAD3A* RNA and ASO were injected separately into fertilized zebrafish eggs prior to the 4-cell stage. Embryos were collected and imaged at 3dpf or 4dpf. When using *atad3*-null zebrafish crossing, embryos were separately collected in 50mM NaOH for DNA extraction and genotyping following imaging. Genotyping was performed using allele-specific primers for both the WT allele and the allele carrying a CRISPR/Cas-9 mediated deletion (primers are listed in **Supplemental Table 1**). Images of zebrafish embryos were acquired using Discovery.V8 stereoscope and AxioCam MRc digital camera (Zeiss) while embryos were mounted on their side in 2% methyl cellulose.

## Data availability statement

Data are available upon request from the authors.

## Acknowledgements

T.Ha. and W.H.Y. are supported by a joint grant from the United States-Israel Binational Science Foundation (BSF 2023188) and the 2024 Neufeld Memorial Research Grant. T.Ha. is also supported by the Israel Science Foundation (ISF 3260/21). W.H.Y. is also supported by the Oklahoma Medical Research Foundation and the Presbyterian Health Foundation (4411-12-13-1). M.B.M. is supported by the GeroScience Training Grant (T32AG052363) of the National Institutes of Health (NIH). T.Hw. is supported by the National Institutes of Health (NIH) grant R35GM160013.

The content is solely the responsibility of the authors and does not necessarily represent the official views of the NIH.

## Author contributions

Ezer, S: Conceptualization, Investigation, Formal Analysis, Writing – original draft

Yanovsky-Dagan, S: Investigation, Formal Analysis, Writing – review and editing

Granit, A: Investigation, Formal Analysis

McDougal, M: Investigation, Formal Analysis, Writing – review and editing

Hwang, T: Investigation, Formal Analysis

Antman, I: Software, Formal Analysis

Karni, R: Supervision, Writing – review and editing

Yoon, WH: Funding Acquisition, Supervision, Writing – review and editing

Saada, A: Investigation, Writing – review and editing

Inbal, A: Resources, Supervision, Writing – review and editing

Harel, T: Conceptualization, Funding Acquisition, Supervision, Writing – original draft

## Declaration of interests

None to declare.

## Notes

### Competing Interest Statement

The authors have declared no competing interest.

