## Supplemental Tables and Figures for "Allele-specific antisense oligonucleotide treatment rescues *atad3-*associated phenotype in zebrafish"

**SUPPLEMENTAL DATA:**

**Supplemental Table 1. Primers used in the study**

**Supplemental Table 2. ASO designs**

**Supplemental Table 3. Number of embryos imaged and measured for each group**

**Supplemental figure 1. Effect of ASO4 derivatives on affected fibroblasts**

**Supplemental figure 2. Exon-skipping additional data**

**Supplemental figure 3. *ATAD3A* RNA sequence for zebrafish injections**

**Supplemental figure 4. WT and not mutant *ATAD3A* rescues *atad3*-null phenotype in 3dpf embryos**

**Supplemental figure 5. Human WT *ATAD3A* effect in zebrafish**

**Supplemental figure 6. Human *ATAD3A* transcripts and their effect in zebrafish, with or without ASO**

### Supplemental Table 1: Primers used in the study

| Purpose | Product | Forward primer | Reverse primer |
|---|---|---|---|
| RT-qPCR (fibroblasts) | Total *ATAD3A* – ex14-16 | GGCCACAGAAGGAAAGCAG | CCGTCCTCGGAGGCATAC |
| | *ATAD3A,* c.1582 T specific | GGCCACAGAAGGAAAGCAG | CTGAGCGATCTCCCA |
| | *ATAD3A,* c.1582 C specific | GGCCACAGAAGGAAAGCAG | TGAGCGATCTCCCGG |
| | *ATAD3A*, Total exon 15 | GGCCACAGAAGGAAAGCAG | AGGACACGGCCAGCTGAG |
| | *ATAD3A,* Exon 15 skipping | CACAGAAGGAAAGCAGCCAC | CCTTCAGCCAGCACATCTTC |
| | *RPLP0* (housekeeping) | GAAACTCTGCATTCTCGCTTCC | GACTCGTTTGTACCCGTTGATG |
| ONT sequencing (iPSCs) | *ATAD3A* | GCAACAAGTTCATGCTGGTCC | CAGCACATCTTCTGCTGGTG |
| Ampliseq primers | *ATAD3A* with ampliseq adapters (adapter sequence in italics) | *TCGTCGGCAGCGTCAGATGTGTATAAGAGACAG* GGCCACAGAAGGAAAGCAG | *GTCTCGTGGGCTCGGAGATGTGTATAAGAGACAG* CCGTCCTCGGAGGCATAC |
| Genotyping of *atad3* deletion in zebrafish | Allele-specific WT *atad3* | CGGATCTGCTCACCCTTCA | CCTTGATCTTGCCCTGGTGCT |
| | Allele-specific deletion *atad3* | CGGATCTGCTCACCCTTCA | TACCTTGATCTTGCCCTGCCCA |
| Site-directed mutagenesis | Site-directed mutagenesis | CATGTCGGGCTGGGAGATCGCTC | CCCTCCGTCAGCCGAG |
| Plasmid confirmation | Gel – plasmid | TGGCAGTACTCCCATTGACG | GTCTGACGGTGCTCGGC |
| | Sequencing of plasmid | ATTTAGGTGACACTATAG | CACATCTTCTGCTGGTGCTG |
| | Sequencing of mutagenesis region | AGCGCCTGGTGAGAATGTAT | CACATCTTCTGCTGGTGCTG |

**Supplemental Table 2: ASO designs**. c.1582C>T reverse complement variant in bold, additional mismatched in red

| | 5'- 5 2'MOE modified bases | 10 DNA bases | 5 2'MOE modified bases -3' |
|---|---|---|---|
| ASO1 | CTCCC | **A**GCCCGACAT | GCCCT |
| ASO2 | TCTCC | C**A**GCCCGACA | TGCCC |
| ASO3 | ATCTC | CC**A**GCCCGAC | ATGCC |
| ASO4 | GATCT | CCC**A**GCCCGA | CATGC |
| ASO5 | CGATC | TCCC**A**GCCCG | ACATG |
| ASO6 | GCGAT | CTCCC**A**GCCC | GACAT |
| ASO7 | AGCGA | TCTCCC**A**GCC | CGACA |
| ASO8 | GAGCG | ATCTCCC**A**GC | CCGAC |
| ASO9 | TGAGC | GATCTCCC**A**G | CCCGA |
| ASO10 | CTGAG | CGATCTCCC**A** | GCCCG |
| ASO4a | GATCT | CCT**A**GCCCGA | CATGC |
| ASO4b | GATCT | CCC**A**ACCCGA | CATGC |
| ASO4c | GATCT | CCC**A**GCCTGA | CATGC |
| ASO4d | GATCT | CCG**A**GCCCGA | CATGC |
| ASO4e | GATCT | CGT**A**GCCCGA | CATGC |
| ASO4f | GATCT | CTT**A**GCCCGA | CATGC |
| Scrambled ASO | GACAT | GCCCCTAGCG | ATCTC |

**Supplemental Table 3: Number of embryos imaged and measured for each group**

| | Injected RNA | Number of embryos imaged and measured |
|---|---|---|
| Fig. 3A,B | Un-injected embryos | 6 |
| | WT RNA, 125pg | 7 |
| | WT RNA, 250pg | 3 |
| | WT RNA, 375pg | 3 |
| | Mutant RNA, 125pg | 14 + 1 under-developed |
| | Mutant RNA, 250pg | 12 + 3 under-developed |
| | Mutant RNA, 375pg | 12 + 6 under-developed |
| Fig. 4C | Un-injected embryos | 28 |
| | Mutant RNA, 250pg | 28 |
| | Mutant RNA, 250pg + ASO4a, 100pg | 23 |
| | Mutant RNA, 250pg + scrambled ASO, 100pg | 23 |
| | ASO4a, 100pg | 20 |
| | Scrambled ASO, 100pg | 21 |

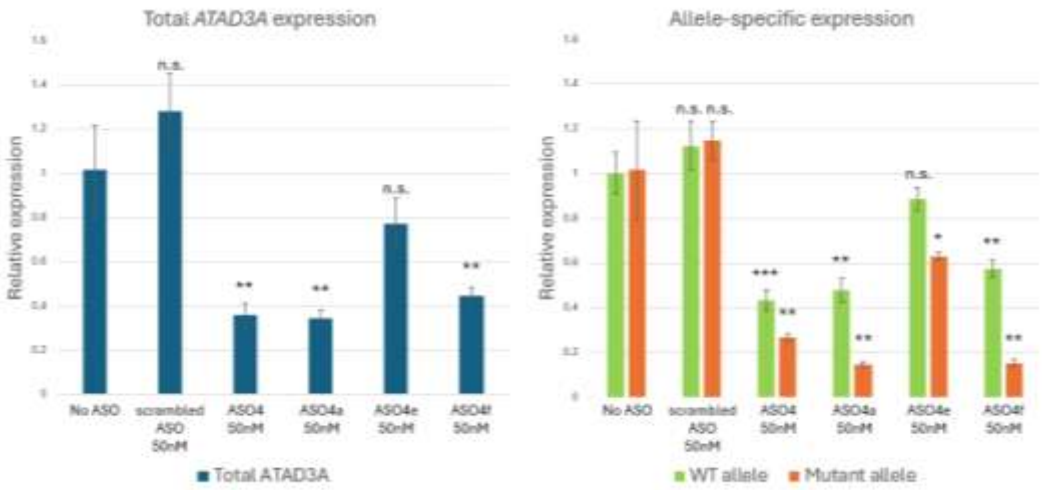


**Supplemental figure 1. Effect of ASO4 derivatives on affected fibroblasts.** Total and allele-specific *ATAD3A* expression in affected fibroblasts treated with 50nM of ASO4, ASO4a,e,f and scrambled ASO compared to untreated cells. Asterisks above each column represent level of significance compared to untreated affected cells (*p val<0.05, **p val<0.01, ***p val<0.001, n.s. not significant).

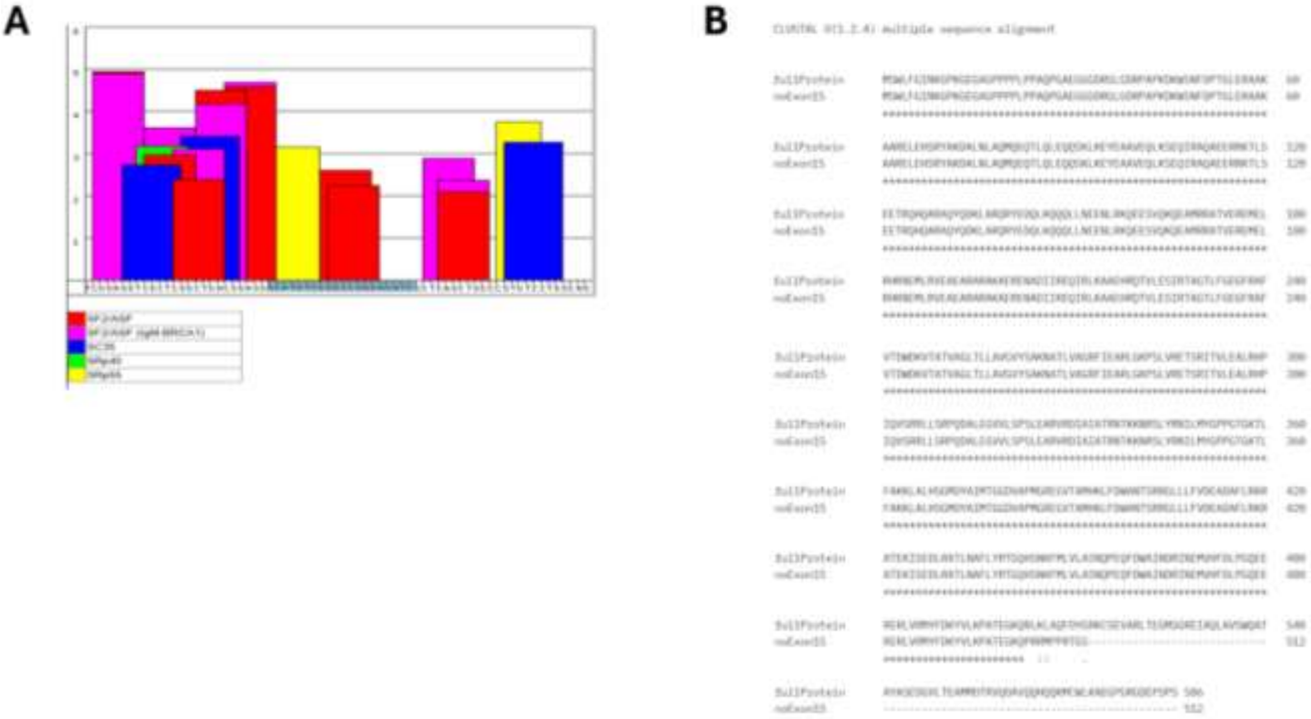


**Supplemental figure 2. Exon-skipping additional data.** (A) ESE-finder results for the *ATAD3A* c.1582 region. In grey is the region complementary for ASO4 and its derivatives, including SF2/ASF and SRp55 binding sites (each color represents a splicing-factor binding site, as indicated in the image from ESE-finder). (B) Clustal Omega alignment of the full ATAD3A protein sequence and the protein lacking exon 15.

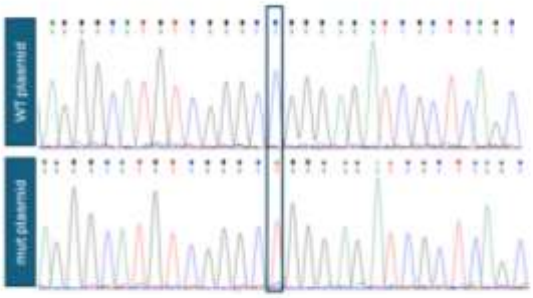


**Supplemental figure 3.** ***ATAD3A* RNA sequence for zebrafish injections.** Sequencing confirming the c.1582 C or T variant (marked by a blue box) in the WT and mutant plasmid, respectively.

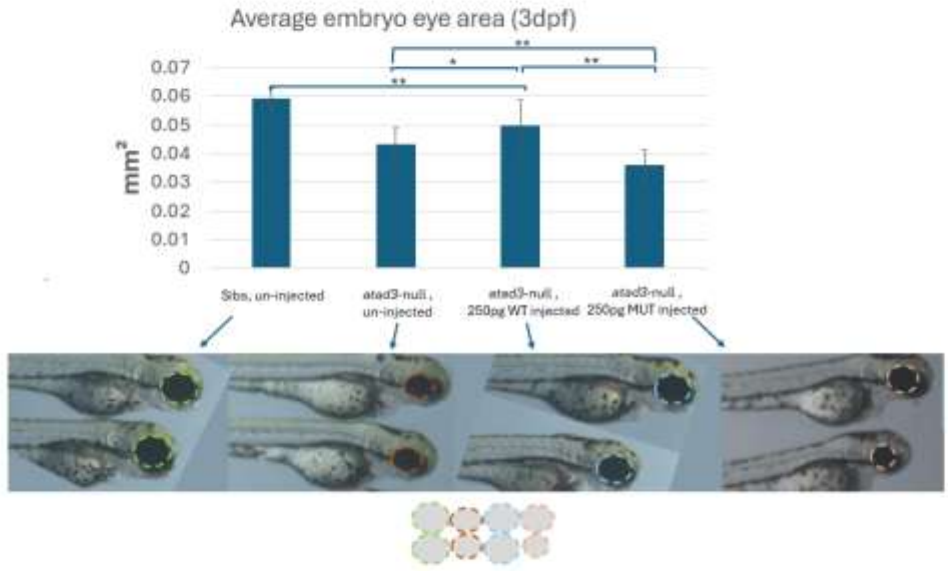


**Supplemental figure 4. WT and not mutant *ATAD3A* rescues *atad3*-null phenotype in 3dpf embryos.** Injection of 250pg WT *ATAD3A* RNA to *atad3*-null embryos increased eye size at 3dpf compared to un-injected *atad3*-null embryos, while injection of 250pg of the mutant human RNA further decreased eye size compared to the un-injected *atad3*-null embryos. Representative images of eyes of 3dpf injected or un-injected embryos are presented.

Numbers of embryos imaged and measured**:** un-injected, sibs: n=13 eyes (13 embryos); un-injected, *atad3*-null: n=11 eyes (11 embryos); 250pg WT, *atad3*-null: n=16 eyes (11 embryos); 250pg mutant, *atad3*-null: n=13 eyes, 7 embryos; 3 additional embryos with extra small eyes could not be measured.

Asterisks represent levels of significance (*p<0.05, **p<0.01).

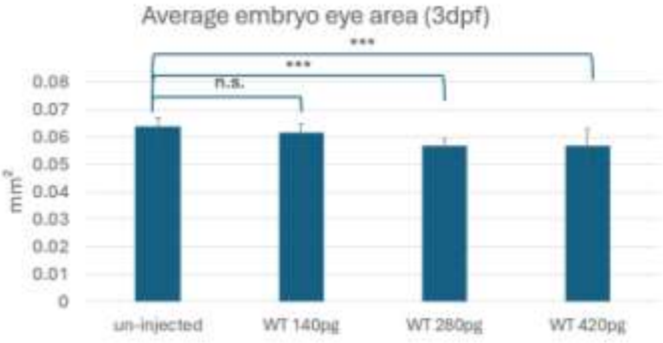


**Supplemental figure 5. Human WT *ATAD3A* effect in zebrafish.** Average eye size at 3dpf of embryos injected with different amounts of WT *ATAD3A* RNA. Embryos images and measured: un-injected n=13; 140pg n=13; 280pg n=5; 420pg n=10. Asterisks represent levels of significance (***p<0.001, n.s. not significant).

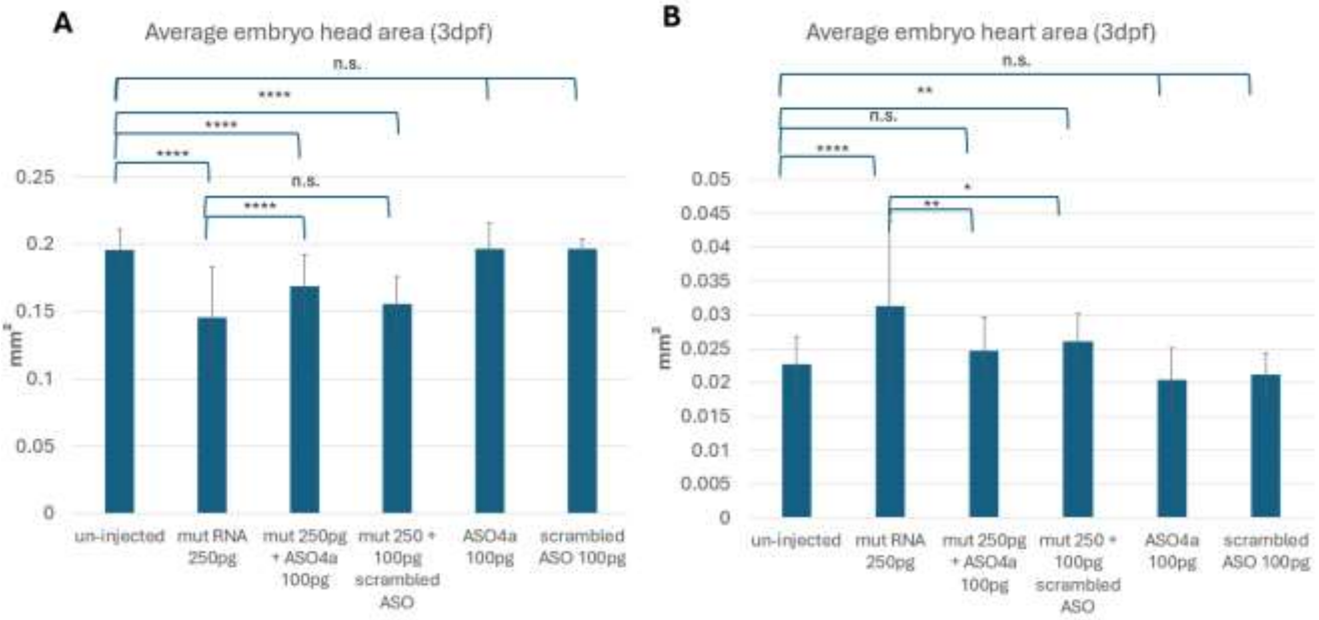


**Supplemental figure 6. Human *ATAD3A* transcripts and their effect in zebrafish, with or without ASO. (A)** Average head size at 3dpf. Embryos are either un-injected on injected with RNA and/or ASO, as indicated. **(B)** Average heart size at 3dpf. Embryos are either un-injected on injected with RNA and/or ASO, as indicated. Asterisks represent levels of significance (*p<0.05, **p<0.01, ***p<0.001, ****p<0.0001, n.s. not significant).
